# Accelerating multi-objective V_H_H discovery via integrated high-throughput selection and AlphaFold3-guided structure prediction

**DOI:** 10.64898/2026.01.19.700436

**Authors:** Max D. Overath, Suthimon Thumtecho, Esperanza Rivera-de-Torre, Melisa Bénard-Valle, Darian S. Wolff, Rahmat Grahadi, Kasper H. Björnsson, Andreas S.H. Rygaard, Nils Hofmann, Jann Ledergerber, Anne Ljungars, Andreas H. Laustsen, Simon Olsson, Thomas J. Fryer, Timothy P. Jenkins

## Abstract

Discovering therapeutic antibodies that bind multiple related targets with high affinity and favourable biophysical properties remains challenging and resource intensive. For snakebite antivenoms, this challenge is critical as treatments must neutralise toxins across multiple snake species. We developed a pipeline combining high-throughput yeast screening, deep sequencing, and AlphaFold3 structure prediction to rapidly identify poly-specific variable domains of heavy-chain-only antibodies (V_H_Hs) against long-chain α-neurotoxins. Multiplexed yeast display screening generated a dataset of diverse candidates with varying binding specificities. AlphaFold3-generated V_H_H-toxin complex predictions enabled structure-based prioritisation that accurately predicted poly-specific binders targeting conserved epitopes across multiple toxins. These structural insights enabled computational optimisation of both affinity and solubility without disrupting target recognition. Experimental validation confirmed improved variants maintained broad specificity across toxins. This integrated approach accelerates multi-objective antibody discovery by predicting which candidates will bind multiple targets before extensive laboratory testing, providing a generalisable strategy applicable beyond antivenoms to any therapeutic requiring broad target coverage.

## Introduction

Therapeutic antibodies have become well-established treatment modalities, widely used to manage and treat a variety of diseases, including cancers, autoimmune disorders, and infectious diseases, by specifically targeting and neutralising disease-associated antigens [1]. Advances in experimental workflows and computational methods, particularly machine learning (ML), have streamlined antibody discovery and development in recent years [2–5]. Despite this progress, antibody discovery and development remains a time- and resource-intensive endeavour. To deliver antibodies for therapeutic use it is key to optimise for affinity and specificity against a desired epitope across a defined target homologue space, whilst also mitigating off-target binding [6]. These criteria often necessitate multiple rounds of selection to yield optimal lead candidates; a time- and resource-intensive process that is especially limiting for complex target spaces in which binding across multiple protein homologues is required for therapeutic efficacy, as in the development of snake antivenom.

Snakebite envenomation is a devastating health burden, which causes an estimated 100,000 deaths annually [7], underscoring the urgent need for improved therapies against this disease. Recombinant antibodies have emerged as a promising alternative to traditional plasma-derived antivenoms [8–11], as it is hypothesized that recombinant antivenoms can be developed to become safer and more efficacious than their traditional counterparts [12,13]. However, the discovery and development of these therapeutics come with several challenges of therapeutic antibody development. To develop poly-specific recombinant antivenoms with broad species coverage, a central objective is to identify broadly neutralising antibodies that retain poly-specific binding to conserved neutralising epitopes on toxin homologues across venoms from multiple snake species [14]. Here poly-specificity denotes breadth of specific binding across toxin variants, whereas broad neutralisation denotes breadth of functional inhibition. Importantly, poly-specific binding to non-neutralising epitopes may not confer protective activity. Beyond broad neutralisation, successful antivenom antibodies should exhibit high solubility and thermal stability, which could facilitate long shelf-life and deployment in remote areas. Lastly, given that snakebite envenomation disproportionately affects low-income regions, cost-effective discovery and production methods are essential to ensure accessibility and affordability [8,15,16].

In this study, we sought to overcome key antibody discovery challenges such as specificity, affinity and solubility by developing a unified, high-throughput pipeline that seamlessly integrates antibody display technologies, deep sequencing, and *in silico* modelling for hit identification and optimisation (**Fig. 1**). We applied this pipeline to the discovery of poly-specific variable domains of heavy-chain-only antibodies (V_H_Hs) against long-chain α-neurotoxins (LNTxs). These highly lethal toxins are typically found in elapids, such as cobras and mambas, and bind and inhibit acetylcholine receptors (AChRs) [17], causing paralysis and muscle failure in victims and prey [18]. They are therefore highly medically relevant targets for the discovery of broadly-neutralising antibodies [10,11,19,20] and other protein binders [21] that can be used to develop recombinant antivenoms. We deliberately chose a V_H_H library for this study, as this antibody format offers distinct advantages over full-length IgGs, such as higher stability, higher neutralisation capacity per mass, and potential for low-cost manufacture, making them especially suitable for use in the context of snakebite envenomation [12,14,22].

**Figure 1:**
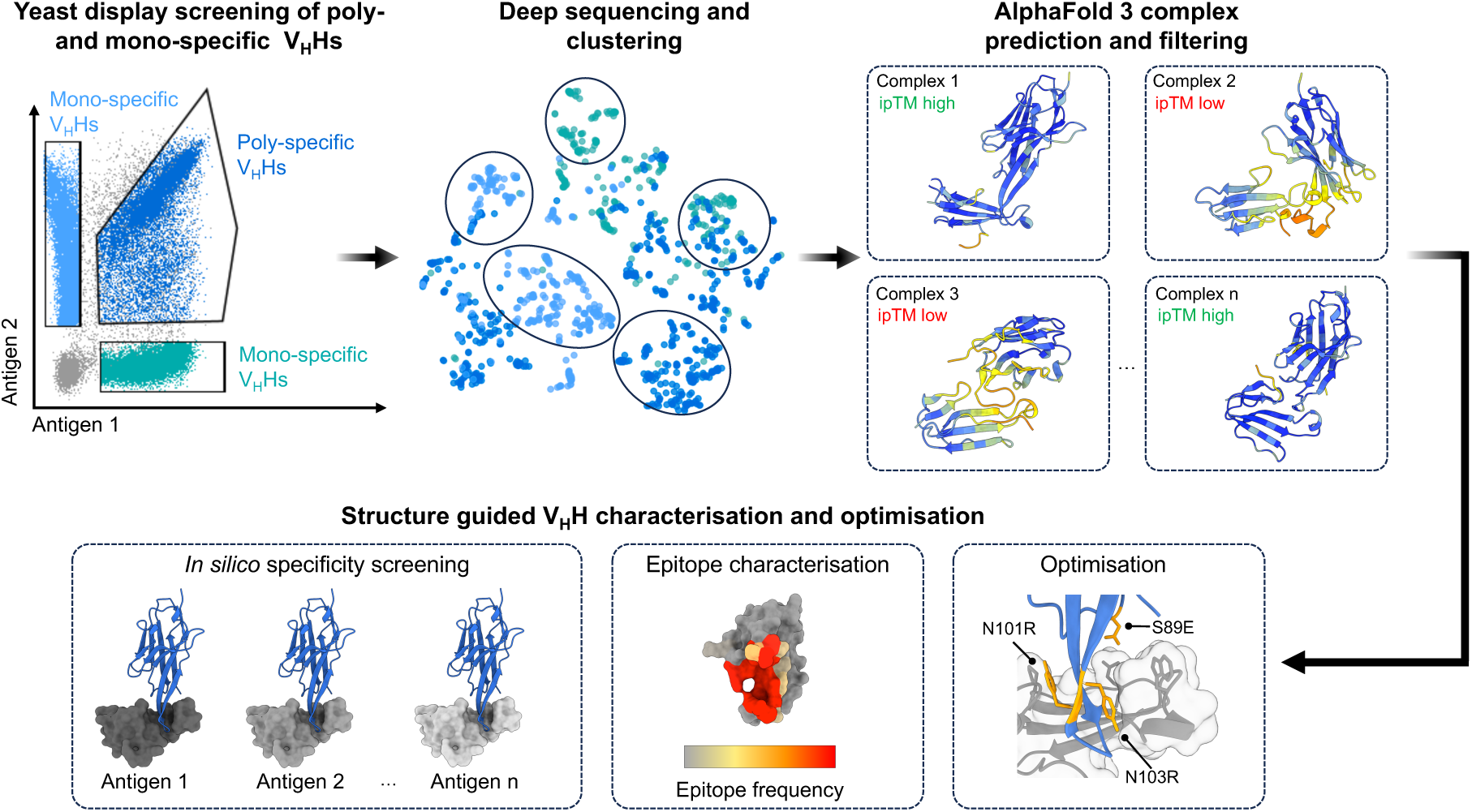
A high-throughput and structure-guided pipeline for the discovery of poly-specific V_H_Hs. Multiplexed yeast display is employed for the high-throughput screening of poly- and mono-specific V_H_Hs against different long-chain α-neurotoxins (LNTxs). The output clone pools are deep sequenced and clustered based on CDR1-3 sequence similarity to downsample and identify relevant V_H_H clones. AF3 is then used to predict complexes of V_H_Hs and respective LNTxs with subsequent scoring of the complexes using the interface predicted template modelling (ipTM) score to select high-confidence predictions. These structures support *in silico* specificity screening and enable structure guided selection, characterisation, and optimisation of poly-specific V_H_Hs.

To enable the discovery of poly-specific V_H_Hs, we developed a streamlined dual-display approach that combined phage and yeast display. The combination of phage and yeast display has previously been shown to effectively integrate the strengths of both techniques [23–25]. Phage display enables efficient selection of binders from large libraries, while yeast display enables quantitative, multiplexed screening capabilities. We used multiplexed yeast-display to efficiently identify poly-specific V_H_Hs binding to distinctly fluorescently labelled toxins. Following the selection and screening campaign, we integrated deep sequencing of all sorted populations (poly-specific, mono-specific, non-binding) to build a comprehensive dataset of all antibody variants. This allows for sequence clustering based on complementarity-determining region (CDR) sequence similarity to reveal the diversity and relationships within the antibody library [26,27]. When applied to both positive and negative selections, as done in this study, deep sequencing further enables improved identification of relevant antibody clones by normalisation based on the enrichment ratios [28]. Collection of both positive and negative labelled sequence data also greatly facilitates the downstream training, validation, or fine-tuning of antibody-specific ML models [29–33].

Recent advances in antibody-antigen structure prediction, particularly with AlphaFold3 (AF3), enable more accurate modelling of biomolecular complexes at scale [34]. Here, we integrate AF3 systematically into a unified, high-throughput antibody discovery pipeline to support structure-guided candidate triage. Predicted complexes allow rapid identification of epitope specificity, prioritisation of candidates targeting putative neutralising regions, and extensive *in silico* screening for poly-specificity across toxin homologues. We then couple these structures to computational optimisation, improving affinity with a structure-informed language model [35] and solubility with CamSol [36], followed by experimental validation using a split-luciferase expression and binding assay. Together, this framework provides a scalable route to discover and optimise V_H_Hs and other antibody therapeutics for snakebite envenomation and other indications.

## Results

### Multiplexed yeast display and deep sequencing enable high-throughput discovery of poly- and mono-specific V_H_Hs against long-chain three-finger α-neurotoxins

We established an experimental pipeline for the high-throughput selection of poly- and mono-specific V_H_Hs, integrating phage bio-panning and multiplexed yeast display. The starting point for this study was an enriched V_H_H library obtained from one round of phage bio-panning against a consensus long-chain three-finger α-neurotoxin (cLNTx), designed as a consensus of multiple long neurotoxins [37]. The initial library was derived from camelid immune libraries generated by immunization with whole snake venoms as described previously [12]. To enable high-throughput screening of poly- and mono-specific V_H_Hs, we shuttled the enriched phage display library to a yeast display platform. Based on this yeast display library, we employed three different fluorescence-activated cell sorting (FACS) strategies using three different toxins namely the cLNTx, α-cobratoxin (αCbtx, UniProt ID: P01391) and α-bungarotoxin (αBgtx, UniProt ID: P60615) as antigens (**Fig. 2a**). Notably αCbtx and αBgtx were both not considered for the design of the cLNTx. However, αCbtx is structurally closely related to the cLNTx (RMSD = 1.25 Å) while αBgtx is more distant to both αCbtx (RMSD = 3.67 Å) and the cLNTx (RMSD = 3.01 Å) (**Fig. 2a**). At the primary sequence level, cLNTx shares 73.24% and 56.94% identity with αCbtx and αBgtx respectively while αCbtx and αBgtx share 54.93% between them. In a first yeast display screening round, we sorted for binding and non-binding V_H_Hs against cLNTx via FACS by using a fluorescently labelled cLNTx to detect binding and SpyCatcher-GFP to detect V_H_H-SpyTag proteins displayed on yeast cells (**Fig. 2b**). This yielded enriched populations for cLNTx binding (cLNTx+/V_H_H display+ from 8.75% to 56.5%) and cLNTx non-binding (cLNTx-/V_H_H display+ from 23.5% to 58%). Next, we employed a poly-specific sort strategy by using two different fluorescent labels for αBgtx and αCbtx (**Fig. 2c**). Here we could enrich three populations: αBgtx mono-binding (αBgtx+/αCbtx- from 0.093% to 31.3%), αCbtx mono-binding (αBgtx-/αCbtx+ from 1.29% to 53.3%) and αBgtx-αCbtx poly-specific (αBgtx+/αCbtx+ from 7.74% to 49.5%). In addition, we sorted the library for mono- and poly-specific V_H_Hs against cLNTx and αCbtx (**Fig. S1a**). This resulted in enriched populations for cLNTx mono-specific binders (cLNTx+/αCbtx- from 1.03% to 56.4%) and cLNTx-αCbtx poly-specific binders (cLNTx+/αCbtx+ from 6.44% to 34.5%). A population of αCbtx mono-specific binders (cLNTx-/αCbtx+) could not be enriched.

**Figure 2:**
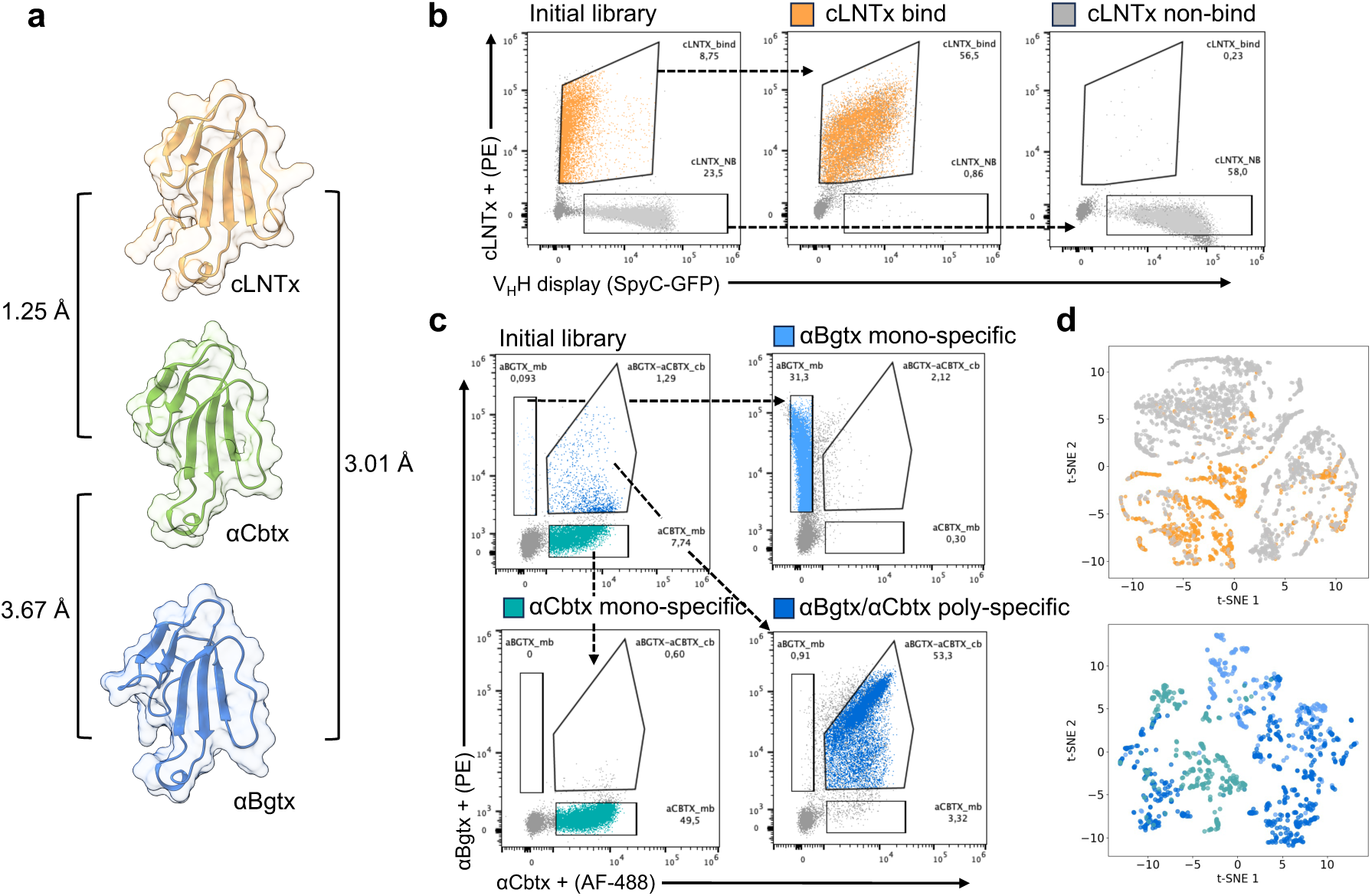
Yeast display guided screening of poly- and mono-specific V_H_Hs. a) Overview of long-chain three-finger neurotoxins (LNTxs) used for the screenings, including a consensus LNTx (cLNTx, structure predicted with AlphaFold3), α-cobratoxin (αCbtx, PDB ID: 1YI5 [38]) and α-bungarotoxin (αBgtx, PDB ID: 1HC9 [39]) with RMSD between structures annotated. b) Yeast display screenings were performed using biotinylated cLNTx with streptavidin-PE to detect antigen binding, and SpyCatcher-GFP to detect V_H_H surface display. Yeast cells were sorted with FACS into cLNTx binder and non-binder populations based on these two signals. c) Poly- and mono-specific selections were performed using biotinylated αBgtx detected with streptavidin-PE and αCbtx directly labelled with AF488. Based on these two signals, three populations were FACS-sorted: αBgtx mono-specific V_H_Hs, αCbtx mono-specific V_H_Hs, and αBgtx/αCbtx poly-specific V_H_Hs. d) Each yeast display selection was deep sequenced, and sequences were clustered by hierarchical clustering using pairwise distances across CDR1-3. Clusters were visualized using t-distributed stochastic neighbour embedding (t-SNE) plots, enabling selection of relevant variants for each sample. Colouring of populations is the same as in (b) and (c).

We then deep-sequenced the enriched yeast display populations. Sequencing reads from paired sorted outputs for each selection (cLNTx binder vs non-binder; cLNTx/αCbtx poly-vs mono-specific; and αBgtx/αCbtx poly- vs mono-specific) were processed using an enrichment-based filter to prioritise sequences supported by the selection. The resulting V_H_H sequences were clustered by hierarchical clustering using pairwise distances computed from global alignments of concatenated CDR1-CDR3 sequences, enabling identification of diverse candidate families for downstream AF3 complex prediction. (**Fig. 2d, Fig. S1b**). Most clusters were dominated by sequences from a single sample (**Fig. S2**), suggesting that the selections enriched distinct CDR1-3 sequence families that may correspond to different binding behaviours. To ensure broad coverage of sequence space, we selected at least two V_H_Hs per cluster by random sampling, ensuring each cluster contributed candidates for AF3 complex predictions.

### AF3 enables high-confidence prediction of multiple V_H_H-toxin complexes

Classically, the next step in antibody discovery is to screen candidates for affinity and specificity, and to map their epitopes to identify the binding site and assess functional relevance. In practice, epitope mapping is typically performed either by solving antibody-antigen complex structures or by mutational scanning approaches such as alanine scanning; both can be resource-intensive and slow and often become a bottleneck when many candidates must be evaluated. Subsequently, lead candidates are engineered to optimise properties such as affinity, poly-specificity, solubility, and other developability-related biophysical parameters. Accurate antibody-antigen structure prediction would greatly streamline these stages. Recent structure prediction models, in particular AF3, have improved on predicting antibody-antigen complexes, but performance remains imperfect, and it has not yet been demonstrated how reliably such predictions can be integrated into an end-to-end antibody discovery and engineering pipeline. To investigate whether AF3 had the ability to overcome these shortcomings and accurately predict high-confidence V_H_H-LNTx complexes, we evaluated AF3 predictive performance on a structure of a V_H_H (TPL1158_01_C09 (C09)) in complex with αCbtx which we had previously described (**Fig. 3a**). C09 originated from the same phage display library used for this study and the complex structure of C09 with αCbtx was resolved by X-ray crystallography (PDB ID: 9GCN) [37]. It is worth mentioning that this structure was released after the AF3 training data cutoff and was not used as a template for predictions. As such it should present an unbiased example of AF3’s ability to accurately predict V_H_H-LNTx complexes. Encouragingly, we found that the predicted complex aligned closely with the crystal structure (RMSD = 0.902 Å, DockQ = 0.813) and showed an identical binding mode of the CDR3 loop (**Fig. 3a**). Since C09 was also shown to be poly-specific against four additional LNTxs, AF3 capabilities to predict this poly-specificity were also benchmarked, as well as differentiating between binding and non-binding using decoy 3FTxs targets (three short-chain α-neurotoxins and three cytotoxins; **Fig. 3b**). To score predicted complexes, we used the interface predicted TM-score (ipTM). We used ipTM > 0.8 as a conservative high-confidence threshold, consistent with published guidance on interpreting ipTM for multimer/complex predictions [40]. This choice is also consistent with prior antibody design studies reporting that higher AF3 ipTM values are enriched among experimentally successful binders and that applying ipTM-based filters can improve hit rates [41,42]. Consistently, every C09:LNTx complex achieved a score above the threshold where expected, whereas all decoy targets exhibited ipTM values well below this threshold, strongly suggesting that AF3 can discriminate between binders and non-binders.

**Figure 3:**
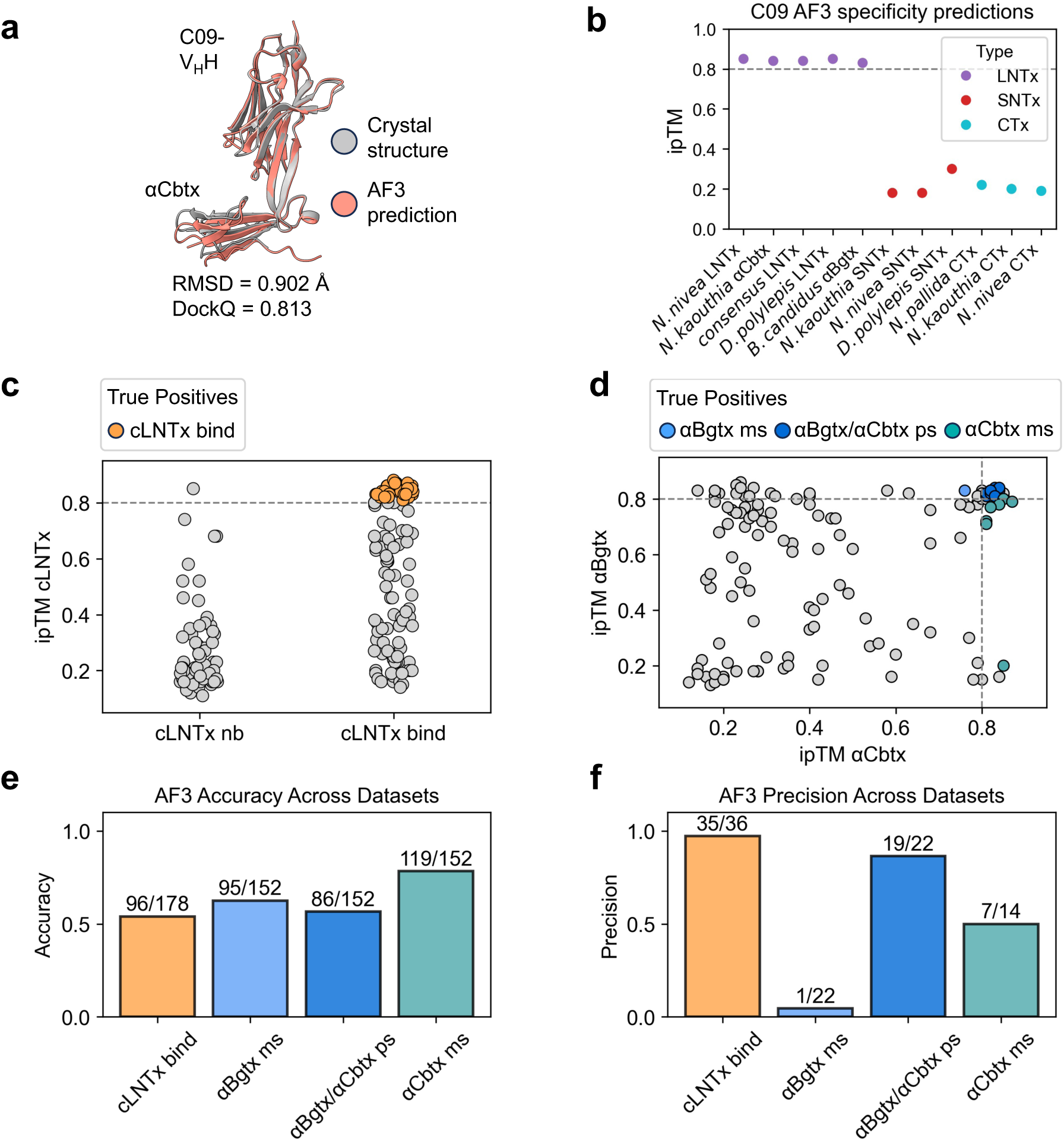
AF3 predictions of V_H_Hs in complex with long-chain α-neurotoxins (LNTxs). a) Accuracy of AF3 prediction is benchmarked using a previously resolved crystal structure of a V_H_H (TPL1158_01_C09 (C09)) in complex with α-cobratoxin (αCbtx) (PDB ID: 9GCN [37]), demonstrating high alignment between predicted and resolved structure (RMSD = 0.902 Å, DockQ = 0.813). b) AF3 enables the prediction of experimentally validated poly-specificity of C09 with four other LNTxs as well as non-binding to decoy targets (three short-chain α-neurotoxins (SNTx) and three cytotoxins (CTx), using the interface predicted template modelling (ipTM) as a scoring parameter (threshold 0.8). c) AF3 was used to co-fold 174 V_H_H sequences from the cLNTx binder and non-binder sets together with the cLNTx sequence. Structures with an ipTM confidence score of at least 0.8 were labelled as predicted binders, and true positives are colour coded. d) From the αBgtx/αCbtx poly-/mono-specific dataset, 152 V_H_H sequences were selected and co-folded with αBgtx and αCbtx sequences in separate AF3 predictions. Predictions were labelled as correct when the predicted binding pattern based for both toxins matched the yeast display selection category for each V_H_H. e-f) Accuracy and precision of predicted mono- and poly-specificity and true experimental binding based on the sequencing data. (nb: non-bind, ps: poly-specific, ms: mono-specific).

To evaluate whether AF3 could accurately predict V_H_H-LNTx complexes and thereby streamline candidate prioritisation, we used AF3 to model structures for V_H_H sequences selected from our clustered deep-sequencing data. As before we utilised an ipTM score > 0.8 as our binding threshold and benchmarked AF3’s predictions against experimental binding data from yeast display screening across three datasets. AF3 demonstrated strong precision in identifying true binders. Across all datasets, 69 of 73 high-confidence predictions (ipTM > 0.8) matched experimental binding labels (precision = 0.94). However, this high precision came at the cost of sensitivity, as many experimentally validated binders fell below the ipTM threshold, resulting in moderate overall accuracy (**Fig. 3c–f**). Notably, AF3 performance varied by binding specificity. For the cLNTx dataset (174 V_H_H sequences), AF3 identified 32 predicted binders, of which 31 were experimentally confirmed (precision = 0.97). For the αBgtx/αCbtx dataset (152 sequences), AF3 excelled at identifying poly-specific binders (19/22 correct, precision = 0.86) but struggled with mono-specific binders, particularly for αBgtx (1/22 correct, precision = 0.04). The cLNTx/αCbtx dataset further confirmed this trend, with perfect precision for poly-specific binders (19/19 = 1.00) but lower precision for mono-specific candidates (9/15 = 0.60) (**Fig. S3**). In total, AF3 generated 77 unique high-confidence predictions from 258 evaluated sequences. Overall, poly-specific predictions achieved a precision of 0.94 (69/73), compared with 0.33 (17/51) for mono-specific predictions. These results demonstrate that AF3 can be effectively integrated into the discovery workflow to generate high-confidence complex structures and prioritise poly-specific V_H_Hs. However, the substantial number of false negatives and the challenges in mono-/poly-specific discrimination indicate that AF3 is best used as a stringent filter for confident binders rather than a comprehensive screening tool.

### AF3 predictions enable epitope characterisation and are experimentally confirmed

Since there was a strong correlation between high-confidence predictions of AF3 and true experimental binding, we reasoned that the predicted high-confidence complexes were accurate enough to analyse epitopes and binding modes across the different selections. One hypothesis we focused on was that toxin epitopes targeted by mono-specific V_H_Hs would likely be distinct from V_H_Hs that are poly-specific against different LNTxs. We thus aligned the AF3-obtained true-positive complexes for each V_H_H selection based on the respective LNTx structure (**Fig. 4a**). The cLNTx bind complexes revealed two distinct clusters of non-overlapping epitopes and three major V_H_H binding modes. The αBgtx/αCbtx poly-specific selection consists of a single non-overlapping epitope shared by all V_H_Hs. Similarly, the αBgtx mono-specific and αCbtx mono-specific V_H_Hs bound to the same non-overlapping epitope, although the binding mode of αCbtx mono-specific V_H_Hs was distinct from that of αBgtx poly-specific V_H_Hs. These findings substantiated our initial hypothesis of distinct mono- vs poly-specific epitopes.

**Figure 4:**
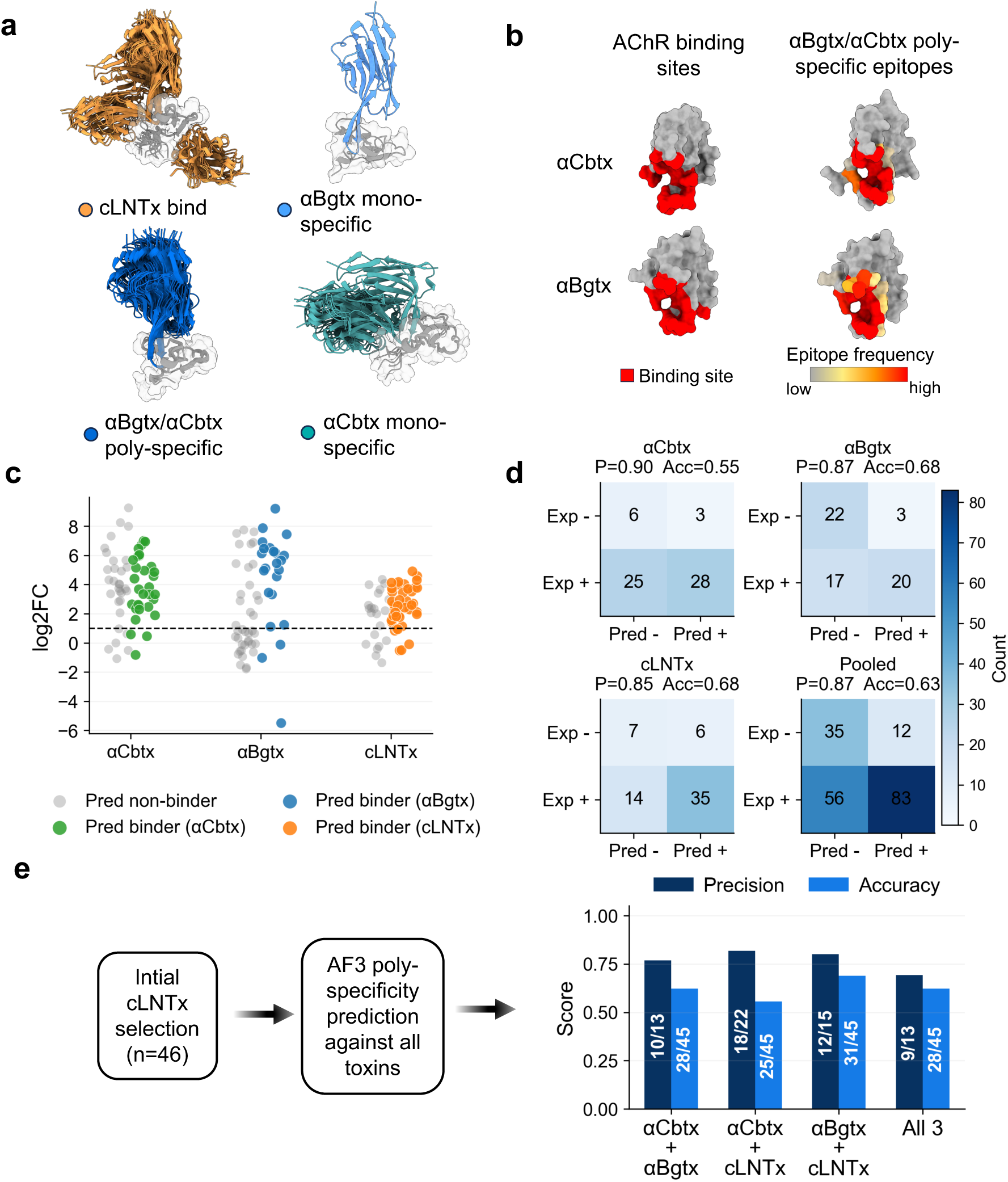
Characterisation and experimental validation of AF3 predicted complexes. a) Representative AF3-predicted V_H_H-toxin complex structures for true-positive predictions (ipTM > 0.8) within each yeast-display sample, i.e., V_H_Hs whose predicted specificity matches the expected population (cLNTx bind, αBgtx mono-specific, αBgtx/αCbtx poly-specific, and αCbtx mono-specific). Toxins are shown in grey. cLNTx bind samples reveal three major V_H_H binding modes that converge on two epitope clusters. The αBgtx mono-specific V_H_H engages an overlapping epitope and adopts a similar binding mode to the αBgtx-αCbtx poly-specific V_H_Hs. αCbtx mono-specific V_H_Hs target a similar epitope region but with a distinct binding mode. b) Comparison of poly-specific V_H_H epitopes with AChR binding sites of LNTxs. Binding sites of αCbtx and αBgtx with the acetylcholine binding protein (AChBP) (PDB ID: 1YI5 [38]) are coloured in red. Epitope residues based on the true-positive αBgtx-αCbtx poly-specific V_H_Hs are coloured based on the epitope frequency and showing strong overlap with AChR binding sites. c) Split-luciferase binding signals (log₂ fold change (log₂FC) over background) for 62 experimentally tested V_H_Hs against cLNTx, αCbtx, and αBgtx. Experimental binding was defined as log₂FC > 1. Points are coloured by AF3 predicted binders (ipTM > 0.8) for the corresponding toxin; grey indicates predicted non-binders. The dashed line marks the binding threshold. d) Confusion matrices comparing AF3 binding predictions to experimental split-luciferase binding for αCbtx, αBgtx, and cLNTx, as well as pooled across toxins. Experimental binding was defined as log₂FC > 1. AF3 predictions were defined as ipTM > 0.8. Cells show counts of V_H_H-toxin pairs; precision (P) and accuracy (Acc) are reported above each matrix. e) Retrospective evaluation of AF3 as a filter for identifying poly-specific binders using only the initial cLNTx selection (binders and non-binders; n=46). AF3 predictions (ipTM > 0.8) were applied against all toxins and compared to experimental split-luciferase binding calls (log₂FC > 1). Performance is summarized as precision and accuracy for V_H_Hs predicted to bind each toxin pair or all three toxins.

The alignment of *in vitro* and *in silico* data thus gave us sufficient confidence to further investigate the predicted structural interactions and actively evaluate whether a given V_H_H would not just bind a target toxin, but would likely also functionally inhibit it. Given LNTxs native mode of action involving binding to and inhibiting the nicotinic acetylcholine receptor (nAChR) we could directly assess whether the V_H_H-toxin epitopes overlapped with the nAChR binding sites. In practice we used a crystal structure of αCbtx bound to acetylcholine-binding protein (AChBP) [38] as a reference and overlaid the poly-specific αCbtx-V_H_H and αBgtx-V_H_H complexes to identify interaction residues (defined as those within 4 Å of the AChBP structure; **Fig. 4b**, left panel). The frequency of each toxin residue being targeted across all poly-specific V_H_Hs was calculated and mapped as a colour gradient onto the LNTx structures (**Fig. 4b**, right panel). This comparison revealed that the epitopes of αCbtx and αBgtx binding V_H_Hs overlap with the receptor binding site, i.e. would have high likelihood to functionally inhibit toxic activity by outcompeting the toxin for receptor binding.

To validate that the obtained high-confidence structures reflect the predicted binding behaviour, we selected a panel of V_H_Hs and experimentally tested them using DNA oligo pool assembly and split-luciferase assays. The panel was designed to probe AF3 performance across outcome classes: true positives (AF3-predicted binders consistent with yeast display selection), representative false positives (AF3 predicted binding but not enriched by yeast display), false negatives (no AF3 predicted binding despite yeast display enrichment), and true negatives (no AF3 predicted binding and originating from the cLNTx non-binding selection). In total, 62 unique V_H_Hs were tested for binding against cLNTx, αCbtx, and αBgtx (**Fig. 4c**). We defined binding as split-luciferase log₂FC > 1 (see Methods and **Fig. S4**) and AF3 ipTM > 0.8, then evaluated predictive performance using confusion matrices per toxin and pooled across all V_H_H-toxin pairs (**Fig. 4d**). Consistent with the trends observed from the yeast display selection, AF3 predictions showed high precision across toxins (αCbtx 0.90, αBgtx 0.87, cLNTx 0.85; pooled 0.87), whereas accuracy was lower (0.55, 0.68, and 0.68; pooled 0.63). These results indicate that AF3-predicted binders were usually correct. However, this high precision comes with a trade-off in sensitivity: a substantial fraction of experimentally positive interactions was not recovered by the ipTM > 0.8 cutoff (pooled false negatives=56), highlighting that AF3 can serve as a stringent filter for confident binders but may miss many true experimental binders under a high-confidence threshold.

Finally, to address the practical question of whether AF3 could have been used up front to enrich for poly-specific binders without additional yeast display screening, we performed a retrospective poly-specificity analysis restricted to the initial cLNTx binder/non-binder selection (n = 46) as a proxy for the initial library after one round of phage panning. Within this set, we evaluated AF3’s ability to identify V_H_Hs predicted to bind toxin pairs for all three toxins (ipTM > 0.8) and benchmarked these predictions against experimental split-luciferase calls (log₂FC > 1), reporting precision and accuracy for each poly-specificity category (**Fig. 4e**). AF3 achieved high precision across categories, i.e. αCbtx+αBgtx, 10/13 (0.77); αCbtx+cLNTx, 18/22 (0.82); αBgtx+cLNTx, 12/15 (0.80); and all three toxins, 9/13 (0.69). Accuracy was again consistently lower, ranging from 0.56 to 0.69 across categories. This suggests that, even without additional screening, AF3 would have provided a high-precision filter for prioritising poly-specific V_H_Hs from the initial selection. Consistent with this, a direct comparison of AF3 and yeast display calls against split-luciferase outcomes showed strong agreement for poly-specific V_H_Hs, whereas mono-specific calls (particularly for αBgtx and cLNTx) were less consistent and often not reproduced by split-luciferase (**Fig. S5)**. Overall, these results reaffirm how AF3 can be incorporated into the discovery process and demonstrate that ipTM values correlate with experimental binding and poly-specificity outcomes.

### Structure guided optimisation of V_H_Hs

While AF3 enabled us to identify likely binding epitopes and select V_H_Hs with high functional potential, the true power lay in its ability to provide a starting point for structural guided refinement of our V_H_Hs with epitope-paratope information. To this end, we employed two previously described methods for antibody optimisation on the 19 high-confidence complexes of αBgtx/αCbtx poly-specific V_H_Hs, since they represented the most relevant V_H_Hs for the potential development of a broadly neutralising antivenom. For solubility optimisation, we applied the CamSol method [36], which proposes mutations predicted to enhance solubility. However, to avoid compromising binding, residues forming the paratope were excluded from mutation. For affinity optimisation, we used a recently developed structure-informed language model [35] that suggests point mutations with potential to improve binding affinity. We therefore designed variants for each V_H_H against both αCbtx and αBgtx separately, retaining only mutations that were predicted to improve binding to both targets, thereby preserving poly-specificity.

Both approaches yielded mutations distributed across CDR and framework regions (FWR) (**Fig. 5a,b**). Notably, mutations for improved affinity were enriched in FWR1 and FWR3, while CamSol mutations were predominantly located in the CDR3 (**Fig. S6**). In total, 43 CamSol-derived and 68 structure-evolution designs were screened using the split-luciferase assay described previously. Several structural evolution (se) variants exhibited markedly enhanced binding signals (up to 4-fold improvement in split-luciferase signal) relative to the wild type for both αCbtx and αBgtx (**Fig. 5c**). Likewise, multiple CamSol variants demonstrated improved expression signals in the HiBiT assay, serving as a proxy for solubility (**Fig. 5d**). Notably, most CamSol variants, while improving expression, did not suffer from significantly reduced binding to αCbtx or αBgtx, potentially due to the structure-informed mutation scheme (**Fig. S7**).

**Figure 5:**
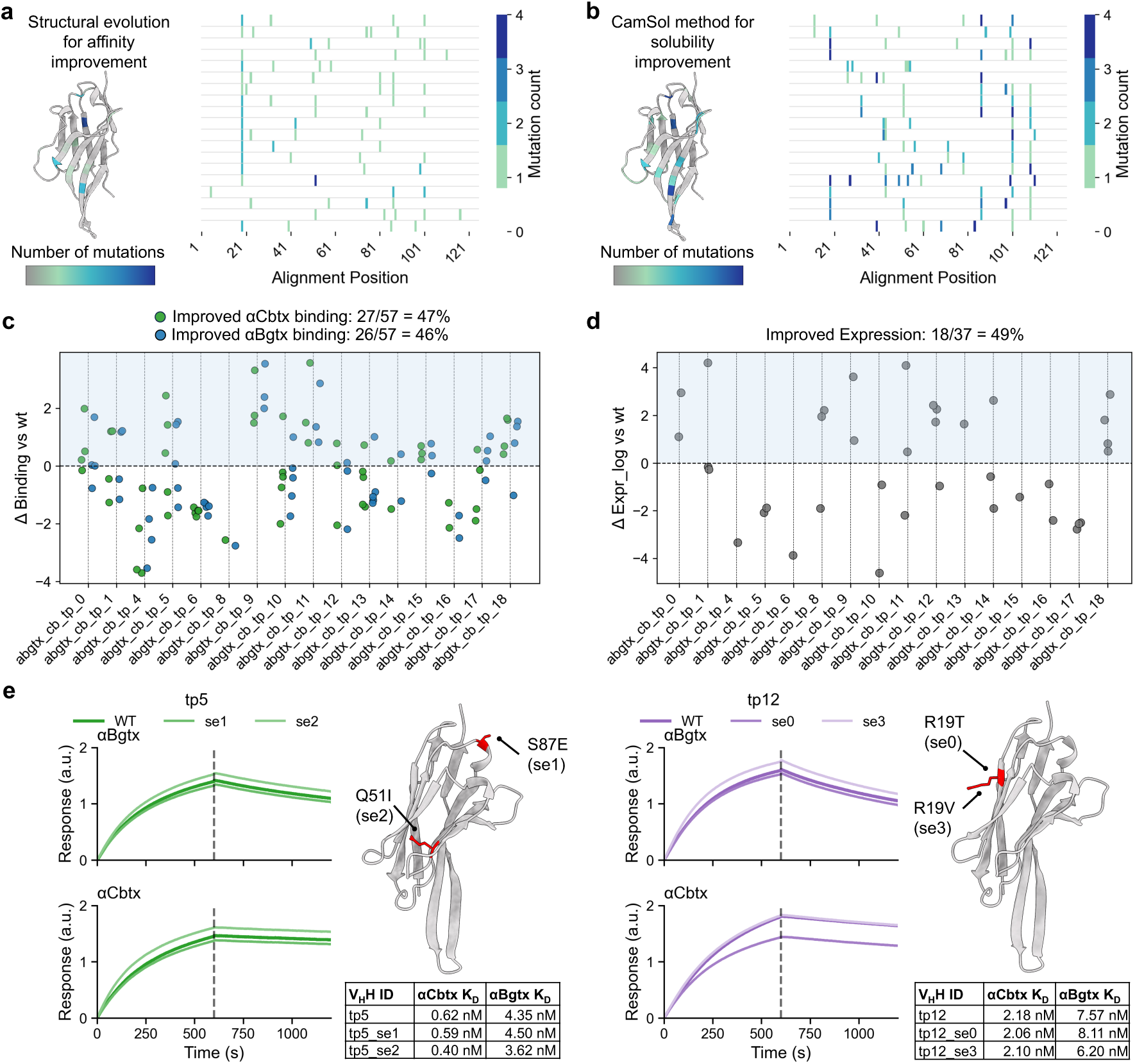
Structure guided affinity and solubility optimisation. a) Mutational landscape of affinity-optimised variants generated from an initial set of 19 wild-type (wt) αBgtx/αCbtx poly-specific V_H_Hs. Each wt V_H_H was diversified using a structural-evolution (se) model, and the resulting substitutions are summarised per alignment position (heatmap; colour indicates mutation count) and mapped onto a reference V_H_H structure to highlight regions preferentially targeted during affinity optimisation. b) Mutational landscape of solubility-optimised variants generated from the same initial set of 19 wt αBgtx/αCbtx poly-specific V_H_Hs. Each wt V_H_H was diversified using CamSol for solubility enhancement, and substitutions are shown per alignment position and projected onto the same reference V_H_H structure to compare the distribution of mutations introduced by solubility-driven optimisation. c) Experimental validation of these variants using the split-luciferase assay, with binding measured against both αBgtx and αCbtx. Shown is the Δ binding of variants vs the wt binding for both αCbtx (green dots) and αBgtx (blue dots). d) Experimental validation of the CamSol variants using the split-luciferase HiBiT assay for expression measurements. Shown is the Δ expression of variants vs the wt e) BLI binding kinetics for tp5 and tp12 WT and two se variants each. Association/dissociation sensorgrams are shown alongside fitted K_D_ values. Structures show the respective V_H_Hs, with mutated residues highlighted in red and each mutation annotated.

To further validate variants with improved affinity, we selected a panel of designed V_H_Hs together with their unmutated wild-type counterparts and measured their affinities for αCbtx and αBgtx using biolayer interferometry (BLI). We first demonstrated that the selected wild-type V_H_Hs already exhibited very high binding affinities for both targets ranging from 7.57 nM to 0.62 nM. (**Fig. S8**). BLI measurements indicated that most variants had affinities similar to, or modestly improved over, the wild type. Several interactions were in the sub-nanomolar regime, where kinetic parameter estimation can approach practical sensitivity limits and should therefore be interpreted cautiously. For example, tp5_se2 showed an apparent improvement against αCbtx (0.62 nM to 0.40 nM) and a modest shift against αBgtx (4.35 nM to 3.62 nM). Likewise, tp12_se3 exhibited a modest improvement against αBgtx (7.57 nM to 6.20 nM) while maintaining comparable affinity to αCbtx (2.10 nM vs 2.18 nM). (**Fig. 5e; Fig. S9**). Notably, all improved variants carried mutations in framework regions distant from the paratope, suggesting that the enhanced affinity may have arisen from improved folding and/or stability of the V_H_H (**Fig. 5e**). Overall, these results demonstrate how structural information on antibody-antigen complexes obtained with AF3 can be directly leveraged to optimize antibody properties.

## Discussion

Rapid, multi-objective antibody discovery pipelines are becoming essential as therapeutic antibody development moves towards broader, faster, and more cost-efficient discovery cycles. Traditional workflows still rely on sequential screening and optimisation, often requiring months to years of iteration to balance affinity, solubility, specificity, and other developability constraints. Here, we show that these objectives can be addressed in parallel by embedding structure prediction and *in silico* optimisation directly into the early discovery phase. We established an integrated pipeline combining multiplexed yeast display, deep sequencing, and AF3-based complex modelling to accelerate discovery and prioritisation of poly-specific V_H_Hs against long-chain α-neurotoxins (LNTxs).

Differentially fluorescently labelled toxins enabled simultaneous selection of mono- and poly-specific V_H_Hs, while deep sequencing provided comprehensive coverage of binder diversity and a benchmark dataset for model evaluation. This combination allowed the identification of poly-specific V_H_Hs at scale and created a foundation for integrating machine-learning models into antibody selection workflows. A key finding of the pipeline was that AF3 can be used as a structural filter within the discovery process rather than purely as a post hoc characterisation tool. In our hands, ipTM > 0.8 yielded a high-precision set of experimentally confirmed binders, consistent with its use as a confidence filter in prior antibody design workflows [41,42]. AF3 therefore distinguished functional binders from non-binders with high precision, particularly for poly-specific V_H_Hs. Our results also suggested that poly-specific candidates can be prioritised directly after an initial enrichment step by using AF3 predictions across multiple toxins, reducing the need for additional experimental specificity screening. Predictive performance was lower for mono-specific clones, reflecting both current limitations of AF3 in modelling antibody-antigen complexes and, in the case of αBgtx and cLNTx mono-specific binders, imperfect initial screening during yeast display. Moreover, the stringent ipTM threshold prioritises precision over sensitivity, and a substantial fraction of experimentally validated binders fell below ipTM greater than 0.8. Increasing the number of sampled complex predictions and adopting ensemble based approaches have been shown to improve multimer prediction [43,44] and are likely to further enhance performance in this setting.

Having access to structural information early in the discovery campaign enables a more rational antibody engineering approach. We used AF3-generated complexes as starting points to guide targeted optimisation of affinity and solubility with a structure-informed language model [35] and the CamSol algorithm [36]. Parallel screening of these variants in a split-luciferase assay rapidly identified V_H_H designs with improved binding signals or expression used as proxy for solubility. Notably, many beneficial substitutions for the structure-informed language fell outside the CDRs, consistent with the structure-informed language model’s premise that conditioning on the full antibody-antigen complex can prioritize mutations that preserve or enhance overall complex stability, including framework changes distal from the interface. Importantly, the V_H_Hs selected for optimisation already exhibited sub-nanomolar to low-nanomolar affinities, underscoring the strength of the upstream discovery workflow while simultaneously limiting the scope for further affinity improvements and making additional optimisation intrinsically more challenging. In this context, the optimisation stage should be viewed primarily as a proof-of-principle that predictive models can be coupled to high-throughput experimental readouts within the same pipeline to tackle multiple objectives in parallel. The pipeline presented here provides a V_H_H-based route that leverages immune-derived diversity, multiplex selection, and structure-guided refinement. Importantly, our approach is agnostic to the specific toxin family and can in principle be applied to other venom components or unrelated antigens where poly-specificity and developability are key design criteria.

Nevertheless, several challenges remain. Some experimentally validated binders did not yield high-confidence AF3 predictions, highlighting current limitations of AF3 for accurate antibody-antigen prediction that may be at least partially mitigated by more extensive conformational sampling and/or explicit modelling of alternative binding modes. While the V_H_H format in combination with small, structurally well-defined toxins is an attractive test case, full-length antibodies and larger, more flexible targets remain difficult for current structure prediction models. In addition, access to AF3 is still limited, although emerging open-source alternatives, such as Boltz-2, Chai-1 and RF3 [45–47], are beginning to close the gap in predictive accuracy and may facilitate broader adoption of similar pipelines.

In summary, this study illustrates how coupling structure prediction with high-throughput selection and screening can help move antibody discovery from a sequential to a more parallelized, predictive process. By integrating multiplexed yeast display, deep sequencing, AF3 complex modelling, and structure-guided optimisation into a single workflow, we provide a scalable foundation for rapid, multi-objective engineering of V_H_Hs and other antibody formats. We anticipate that similar pipelines will be useful not only for recombinant antivenom development, but also for therapeutic, diagnostic, and biotechnological applications where rapid discovery of binders with tailored functional and developability profiles are required.

## Materials and Methods

### Toxins

The consensus long-neurotoxin (cLNTx) was produced in *K. phaffii* KM71H as previously described [37]. Lyophilized forms of α-Cobratoxin (αCbtx; L8114) and α-Bungarotoxin (αBgtx; L8115) were obtained from Latoxan SAS (Portes-lès-Valence, France) and re-suspended in PBS with, and quantified by absorbance at 280 nm (A280). Biotinylation of toxins was done using the Innolink™ Biotin 354S kit (Sigma-Aldrich, Germany) and fluorescent labelling of αCbtx was done using the Alexa Fluor™ 488 NHS Ester Kit (Invitrogen™ A20000) following the manufacturer’s instructions.

### Cloning of phage display outputs into a yeast display vector

V_H_H genes in phagemids from a previously described phage display biopanning against cLNTx [37] were amplified by polymerase chain reaction (PCR) (Q5® High-Fidelity 2X Master Mix, M0492S, New England Biolabs) introducing 30 base-pair overhangs for *in vivo* assembly in yeast by homologous recombination. Primers were:

Homology_fwd:

5′-GGCGGGTCTGGTGGGGGAGGATCTGCCATGCAGGTGCAGCTGCAGGAG-3′

Homology_rev:

5′-TCGCTCGTACCTGGGGTTTCGCTAGCGGCCGCTGAGGAG-3′.

PCR products were purified using the GeneJET PCR Purification Kit (K0702, Thermo Fisher Scientific) following the manufacturer’s instructions. An in-house vector for yeast display containing a SpyTag, N-Terminal Aga2 (Galactose inducible) was digested using NcoI-HF® (R3193L, New England Biolabs) and NotI-HF® (R3189L, New England Biolabs), gel excised and purified using the GeneJET Gel Extraction Kit (K0692, Thermo Fisher Scientific). PCR products and digested vectors were transformed into Saccharomyces cerevisiae EBY-100 strain by chemical transformation as previously described [48].

### Production and purification of SpyCatcher-GFP

A SpyCatcher [49] - GFP fusion protein was recombinantly expressed in *Escherichia coli* (*E. coli*) BL21-(DE3) cells. The SpyCatcher-GFP sequence was encoded on the pET-SpyC-GFP plasmid including a N-terminal His_6_ tag and introduced into cells via chemical transformation. Expression was initiated by diluting 1 mL of a saturated overnight culture into 125 mL of 2xYT medium, followed by incubation at 37 °C with shaking at 250 rpm for 2 hours. Induction was achieved by adding IPTG to a final concentration of 1 mM, and the culture was grown overnight at 25 °C with shaking at 250 rpm. The cells were harvested the next day by centrifugation at 4,800xg for 20 minutes, and the resulting pellet was resuspended in 12.5 mL of equilibration/wash buffer (20 mM sodium phosphate, 500 mM NaCl, 30 mM imidazole, pH 7.4). Cell lysis was performed using sonication, and the lysate was clarified by centrifugation at 4,800xg for 20 minutes. The soluble fraction was used for purification using immobilized metal affinity chromatography (IMAC) with a Ni-NTA resin. Following sample loading, the column was washed with an equilibration/wash buffer to remove non-specifically bound proteins. The target protein was eluted using equilibration/wash buffer + 500 mM imidazole.

### Yeast display selections

Transformed yeast cells were cultured in a selective yeast nitrogen base (YNB)-casamino acids (CAA) medium containing 2% (w/v) glucose at 220 rpm and 30 °C for 12-16 hours. For the induction of V_H_H display, the medium was exchanged to YNB-CAA medium containing 2% (w/v) galactose with an initial cell OD_600_ of 0.5. Cells were incubated for 18–24 hours at 220 rpm and 30 °C. To prepare yeast cells for flow cytometry, cultures were centrifuged at 8,000×g for 1 minute. The cells were washed once by resuspending the pellet in PBSA (PBS with 1 g/L bovine serum albumin (BSA)), followed by another centrifugation step. Biotinylated or AF-488 labelled toxins were diluted to 100 nM in PBSA, and yeast cells were resuspended at 100 μL of toxin solution per 1 OD₆₀₀ unit (i.e., 100 μL/OD). For poly-specific binding sorts, 100 nM of each toxin was used. The cells were incubated with the toxins for 30 minutes at 4°C with shaking at 500 rpm. After incubation, the cells were washed with PBSA and resuspended in a staining mix containing either 1 μg/mL of Streptavidin-Phycoerythrin (PE) (405204, BioLegend) or 500 nM GFP-SpyCatcher. The cells were incubated under the same conditions as before, washed, and resuspended at 1 mL PBSA per 1 OD₆₀₀ unit for fluorescence-activated cell sorting (FACS). FACS was performed using a Sony MA900. After sorting, cells were transferred to SD-UT media and grown for 1-2 days until saturation. Sorting was repeated 2–3 times to achieve enriched populations.

### Illumina deep sequencing

Plasmids from yeast display selections were isolated using the Zymoprep Yeast Plasmid Miniprep II (D2004, Zymo Research). PCR 1 was performed using primers with Nextera adapters and the KAPA HiFi HotStart ReadyMix PCR Kit (KK2601, Roche). Primers were: ngs_pcr1_fwd:

5’-TCGTCGGCAGCGTCAGATGTGTATAAGAGACAGCAGGTGCAGCTGCAGGAG-3’;

ngs_pcr1_rev_2:

5’-GTCTCGTGGGCTCGGAGATGTGTATAAGAGACAGAGCGGCCGCTGAGGAG-

3’.

PCR products were purified using SPRIselect beads (B23318, Beckman Coulter). PCR 2 was performed using Nextera XT Index Kit v2 Set D (FC-131-2004, Illumina) and PCR products were purified again using SPRIselect beads. Quality control was performed using the Qubit™ dsDNA HS Assay Kit (Q32854, Thermo Fisher Scientific) and the Agilent High Sensitivity DNA Kit (5067-4626, Agilent). Sequencing was done using the MiSeq Reagent Kit v2, 500 Cycles (MS-102-2003, Illumina) for paired-end sequencing and spiked with 35% PhiX.

### Processing and clustering of sequencing data

Raw sequencing reads were processed using the MiXCR software package [50] with the following settings: paired-end BCR amplicon analysis was performed using mixcr analyze generic-bcr-amplicon in DNA mode (--dna) with the Lama glama reference (--species lamaGlama). Rigid left and right alignment boundaries were enforced (--rigid-left-alignment-boundary, --rigid-right-alignment-boundary), and clonotypes were assembled across the full V-region span from FR1Begin to FR4End (--assemble-clonotypes-by “{FR1Begin:FR4End}”). To prioritize sequences supported by each selection, we performed an enrichment-based downselection within paired sorted populations for each screening strategy (cLNTx binder vs non-binder; cLNTx/αCbtx poly- vs mono-specific; αBgtx/αCbtx poly- vs mono-specific). For each pair, we computed each sequence’s relative read fraction within each sample and retained sequences whose fraction was higher in the population of interest than in the paired comparator (i.e., positively enriched), discarding sequences depleted in that comparison. For each selection output, retained sequences were clustered using hierarchical clustering based on pairwise distances computed from global alignments of concatenated CDR1–CDR3 sequences. Duplicate CDR1–CDR3 sequences were removed within each sample prior to distance calculation. Hierarchical clustering was performed using Ward linkage, and clusters were defined by cutting the dendrogram at a fixed distance threshold. Dendrogram cut thresholds were set empirically to obtain a tractable number of clusters for downstream analysis: 35 for cLNTx bind/non-bind, 15 for cLNTx/αCbtx poly-/mono-specific, and 10 for αBgtx/αCbtx poly-/mono-specific. Two-dimensional visualisations were generated using t-distributed stochastic neighbour embedding (t-SNE) with n_components = 2, perplexity = 30, and n_iter = 300, applied to the same distance matrix.

### AlphaFold3 prediction and evaluation

V_H_H-toxin complex predictions were performed using AlphaFold3 (AF3) implementation [34] on the AlphaFold Server (https://alphafoldserver.com) using one seed. For each prediction, the best-ranked model was selected, and the ipTM value was extracted. Data were labelled as “predicted binding” if the ipTM value exceeded 0.8. For benchmarking against yeast-display selections, “true binding” labels were derived from the corresponding FACS-sorted populations (binder vs non-binder; mono- vs poly-specific). For experimental validation, “true binding” was defined as split-luciferase log₂FC > 1. Epitopes were defined as antigen residues within a distance of < 4 Å to the V_H_H structure. Epitope analysis was conducted using custom python scripts with biopython/1.84. Complex structures were visualized using ChimeraX-1.6.1.

### Structural evolution

A structure-informed language model [35] was used to propose single-point mutations to potentially improve the affinity of the 19 αBgtx poly-specific V_H_Hs for which high-confidence structures were available. For each V_H_H, the V_H_H-αCbtx and V_H_H-αBgtx complex structures were provided as input to the model. Proposed mutations were then filtered to retain only those suggested for both complexes for a given V_H_H, to prioritize variants that enhance binding without disrupting poly-specificity.

### CamSol

The CamSol method [36] was used via the online server (https://www-cohsoftware.ch.cam.ac.uk/index.php/camsolcombination) to propose solubility-improving mutations for the 19 αBgtx poly-specific V_H_Hs with high-confidence structures. To avoid compromising antigen recognition, residues belonging to the paratope were excluded from the set of positions eligible for mutation, restricting design to non-paratope sites expected to improve solubility without directly perturbing the binding interface.

### Split-luciferase screening of binding and expression

The genes encoding a subset of the discovered V_H_Hs and optimised variants was ordered from Twist as OligoPools and assembled by Golden Gate cloning using BsaI sites. Each V_H_H was fused C-terminally to LargeBiT luciferase, enabling rapid assessment of both expression and antigen binding via a split-luciferase assay (NanoBiT/HiBiT format), conceptually similar to prior implementations [41]. Constructs were transformed into SHuffle® T7 Express *E. coli* T7 cells (C3029J, NEB) and expressed by 1 mM IPTG induction. Cells were lysed and split-luciferase readouts were performed directly on 1:400 diluted lysates. For expression quantification, lysates were incubated with HiBiT detection reagent for 1 h, followed by addition of the luciferase substrate furimazine and measurement of luminescence on a plate reader. For binding measurements, lysates were incubated with biotinylated target antigen for 1 h, then incubated for an additional 1 h with Lumit® streptavidin-SmBiT (W1671, Promega) followed by substrate addition and luminescence measurement. Binding was called positive when the split-luciferase binding readout exceeded background by log₂FC > 1, where background was defined as the luminescence measured for the same V_H_H–LargeBiT lysate incubated without antigen and without SmBiT (V_H_H–LargeBiT + substrate only).

### Expression and purification of selected V_H_Hs

A selection of V_H_Hs was expressed in small-scale cultures for subsequent purification. Expression was initiated by diluting 40 µL of a saturated overnight culture into 4 mL of 2xYT medium, followed by incubation at 37 °C with shaking at 250 rpm for 2 h. Induction was achieved by adding IPTG to a final concentration of 1 mM, and cultures were grown overnight at 25 °C with shaking at 250 rpm. Cells were harvested by centrifugation at 4,800×g for 20 min, and pellets were resuspended in 400 µL PBS (pH 8.0). Cells were lysed by sonication and clarified by centrifugation at 4,800xg for 20 min. The soluble fraction was purified using HisPur™ Ni-NTA Magnetic Beads (88832, Thermo Fisher Scientific) according to the manufacturer’s workflow using PBS (pH 8.0) as equilibration/wash buffer and PBS (pH 8.0) supplemented with 200 mM imidazole for elution. Dialysis to remove imidazole was performed using 3.5 kD MWCO Pierce™ Microdialysis Plates (A50467, Thermo Fisher Scientific).

### Binding analysis using biolayer interferometry

Binding kinetics between purified V_H_Hs and their cognate toxin targets were measured by biolayer interferometry (BLI) using an Octet RED96 instrument (ForteBio) at 30 °C. All measurements were performed in HEPES buffer (pH 7.2). Biotinylated αBgtx and αCbtx (200 nM final) were immobilized on streptavidin (SA) biosensors (Sartorius) to a loading response of ∼1.0 nm. To identify an appropriate analyte concentration for the wild-type (wt) V_H_H, toxin-loaded biosensors were exposed to a 4-point dilution series of V_H_H (300, 100, 33, and 11 nM). For subsequent measurements comparing wt and optimised variants, the V_H_H concentration was fixed at 33 nM. Association was monitored for 600 s followed by dissociation for 600 s in buffer. Data were reference-subtracted using a toxin-loaded sensor incubated without V_H_H. Curves were analyzed in Octet Analysis Studio v12.2.2.26 (ForteBio) and with custom Python scripts.

## Acknowledgments

We thank Irina Borodina (DTU Biosustain) for providing the EBY-100 strain. We also acknowledge Keyan Liu and Viji Kandasamy (DTU Biosustain) for support with the Illumina deep sequencing. Additionally, we thank the HPC support team at the DTU Computing Center (DCC) for valuable assistance in setting up the required computational infrastructure for this work.

## Author contributions

Conceptualization: TPJ, TJF, MDO

Methodology: MDO, TJF, ER, MB, KHB, TPJ

Investigation: MDO, TJF, ST, DSW, RG, ASHR, NH, TPJ

Visualization: MDO, JL

Supervision: TPJ, TJF, SO, AHL, AL

Writing-original draft: MDO, TPJ

Writing-review & editing: All authors

## Data availability

AlphaFold 3 prediction output files for all predicted complexes, together with processed sequencing data, AF3 prediction summary tables, and split-luciferase raw and processed data, are available on Zenodo: https://doi.org/10.5281/zenodo.18290348

---

**Fig. S1.**
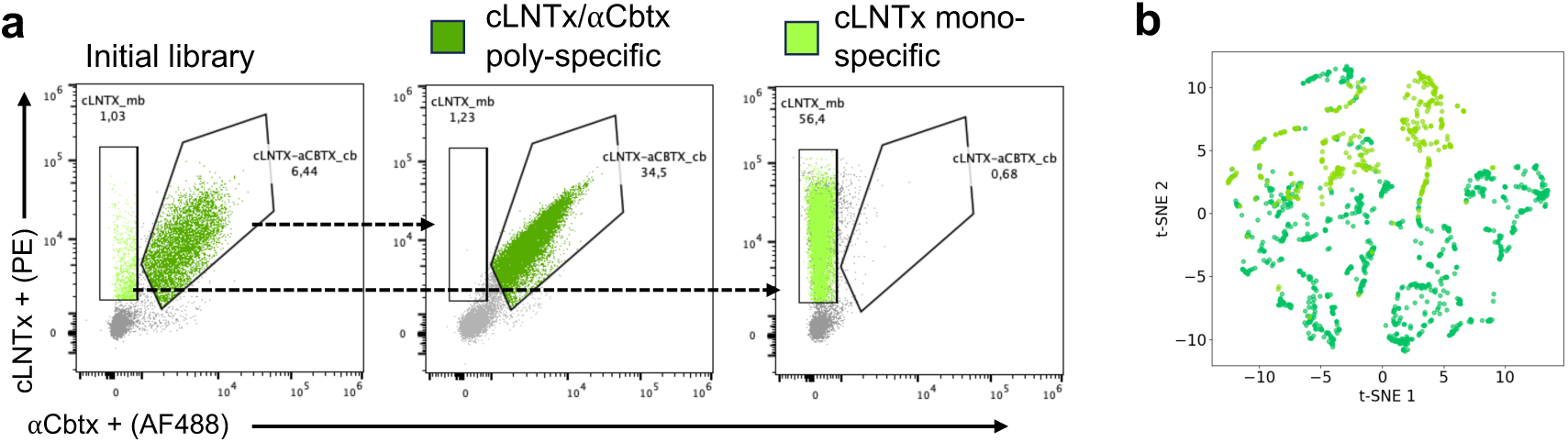
cLNTx/αCbtx yeast display selections and CDR clustering. a) Poly- and mono-specific yeast-display selections using fluorescently labelled cLNTx and αCBTx as antigens. b) Clustering of CDR sequences for cLNTx/αCBTx poly-specific and cLNTx mono-specific samples.

**Fig. S2.**
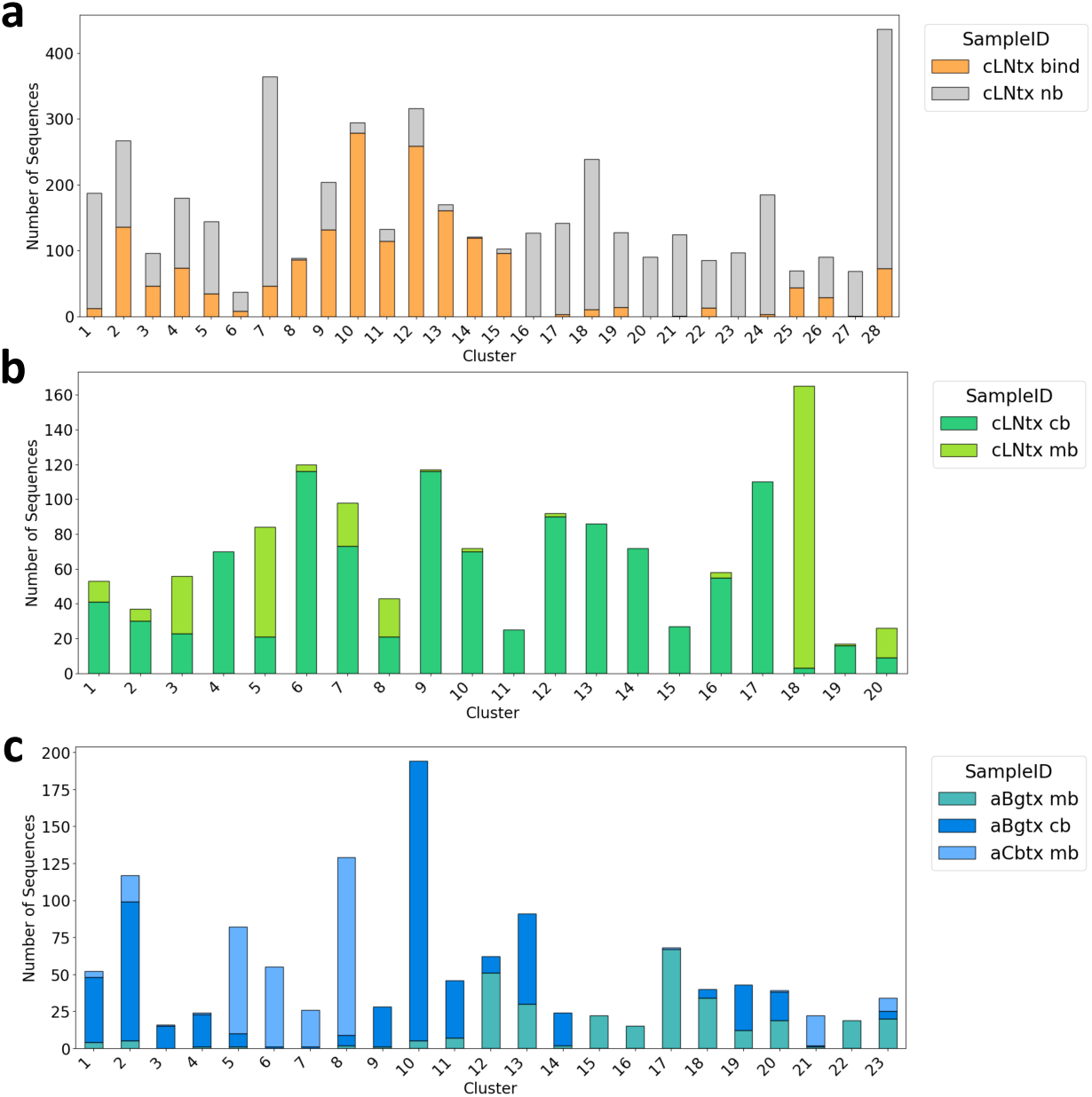
Distribution of different samples per cluster. CDR sequences were clustered using hierarchical clustering of CDR1-3. The plot shows the frequency of each sample within each cluster, revealing that most clusters consist mainly of a single sample. Data were separated by selection for: a) cLNTx bind and non-bind; b) cLNTx/αCbtx poly-specific and αCbtx mono-specific; and c) αBgtx/αCbtx poly-specific, αCbtx mono-specific, and αBgtx mono-specific.

**Fig. S3.**
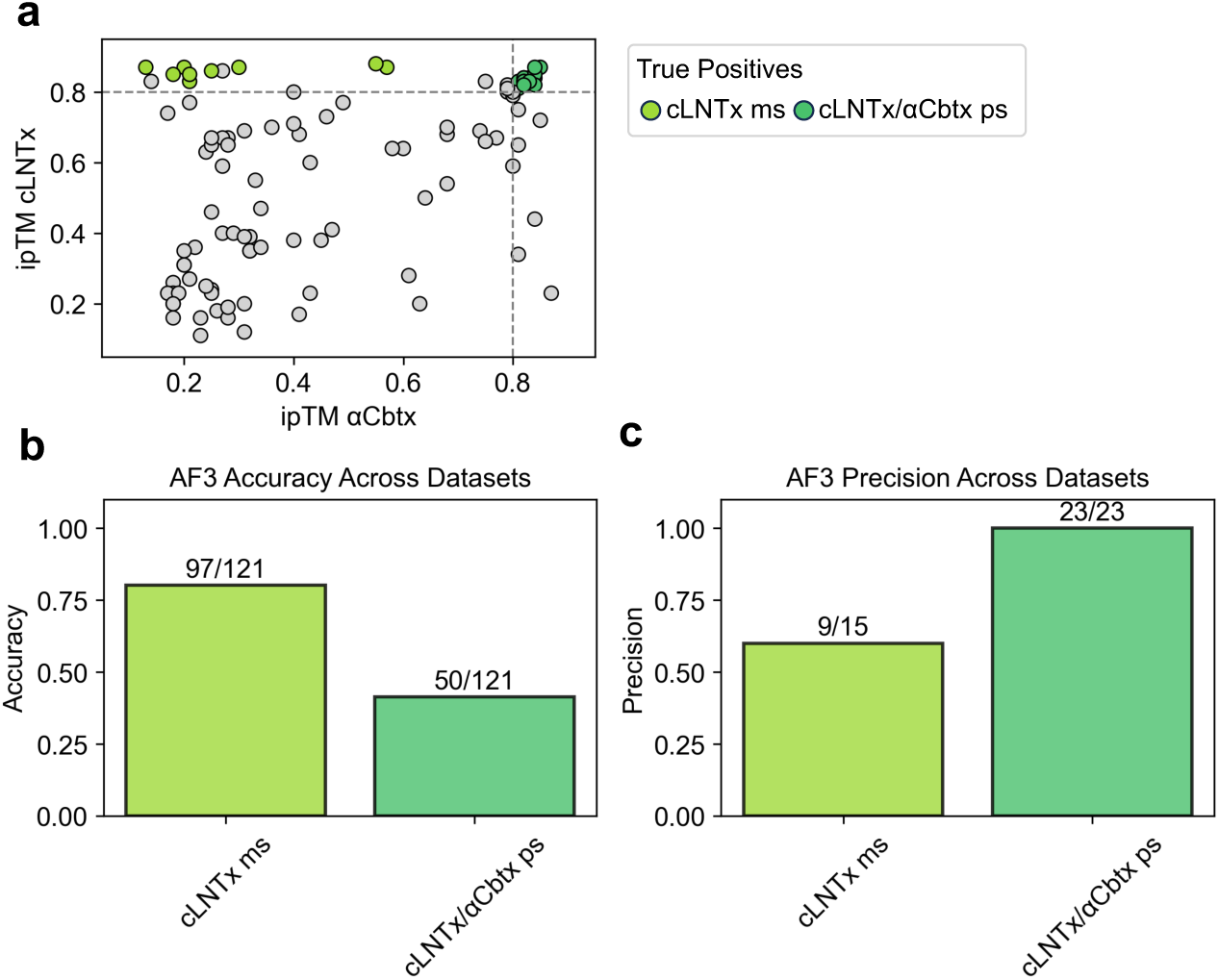
cLNTx/αCbtx AF3 predictions evaluation. a) ipTM distributions for V_H_H-toxin predictions, where colored dots indicate true-positive predictions for cLNTX/αCBTx poly-specific and αCbtx mono-specific samples. b) Accuracy and c) precision plots show very high precision for cLNTX/αCBTx poly-specific V_H_Hs (precision = 1.0) and lower precision for cLNTx mono-specific samples (precision = 0.6), while accuracy is higher for mono-binding samples (accuracy = 0.80) than for cross-binding samples (accuracy = 0.46).

**Fig. S4.**
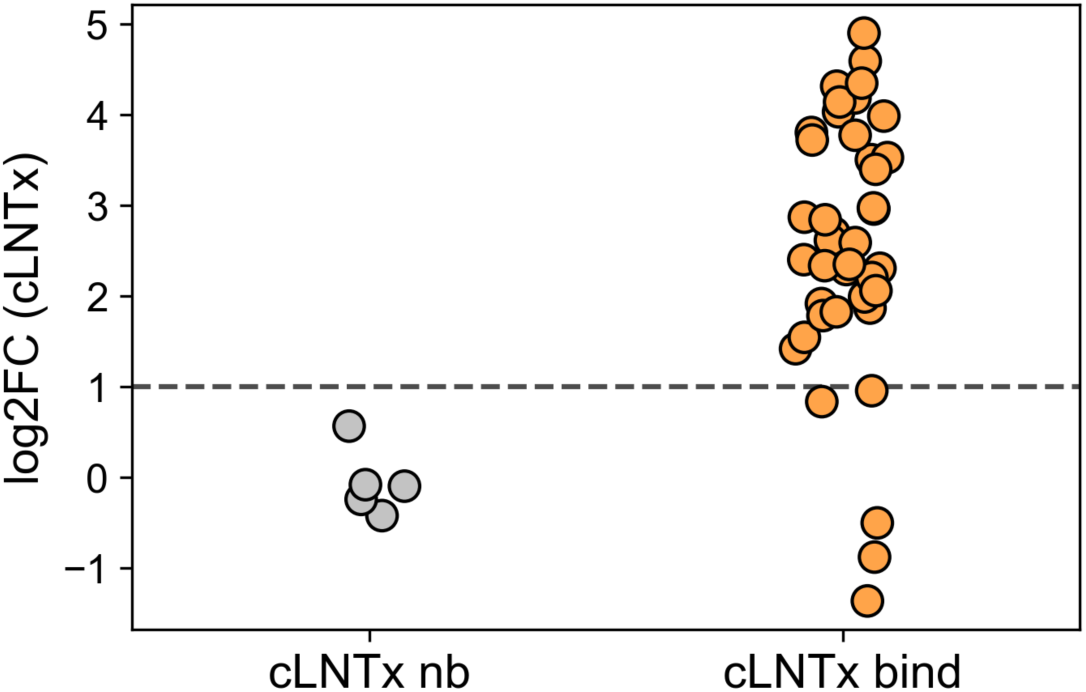
Split-luciferase cLNTx binding cutoff selection. Split-luciferase binding signals (log₂FC) for V_H_H-LargeBiT lysates measured with biotinylated cLNTx and streptavidin-SmBiT, normalized to background (V_H_H-LargeBiT + substrate only). Points are colored by yeast-display label (cLNTx non-binding, grey; cLNTx binding, orange). The dashed line marks the threshold used in this study (log₂FC = 1), which separates the non-binding distribution from the majority of binders.

**Fig. S5.**
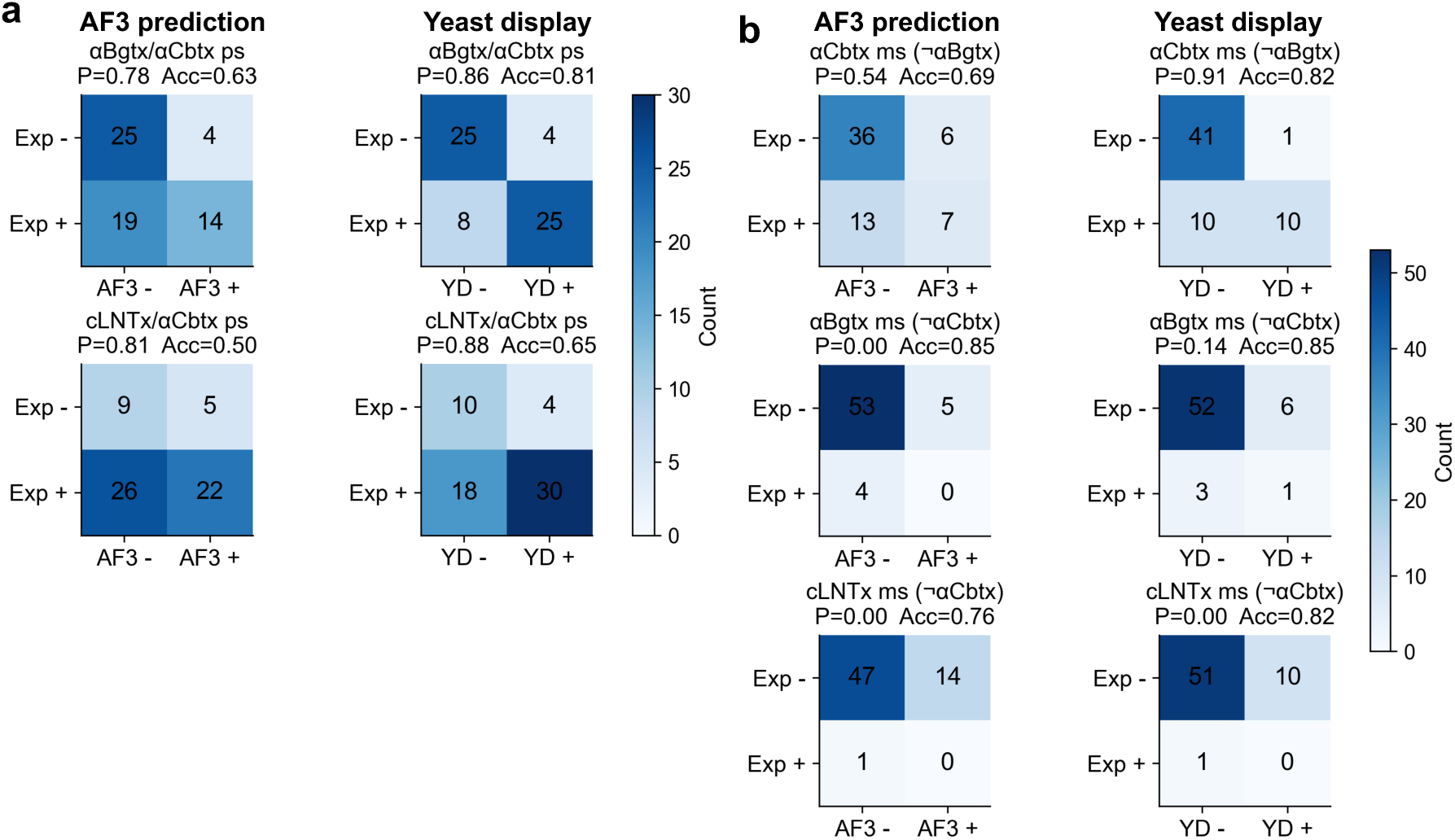
Comparison of AF3 and yeast display (YD) for predicting poly- and mono-specific binding. Confusion matrices benchmark AF3 predictions (ipTM > 0.8; AF+/AF3−) and yeast display (YD) selection calls (YD+/YD−) against experimental split-luciferase binding (Exp+/Exp−; log₂FC > 1). a) Poly-specific (ps) classification for αBgtx/αCbtx and cLNTx/αCbtx V_H_Hs. b) Mono-specific (ms) classification for αCbtx, αBgtx, and cLNTx V_H_Hs (¬ indicates the target that should not be bound). Precision (P) and accuracy (Acc) are shown for each panel; numbers in cells indicate counts.

**Fig. S6.**
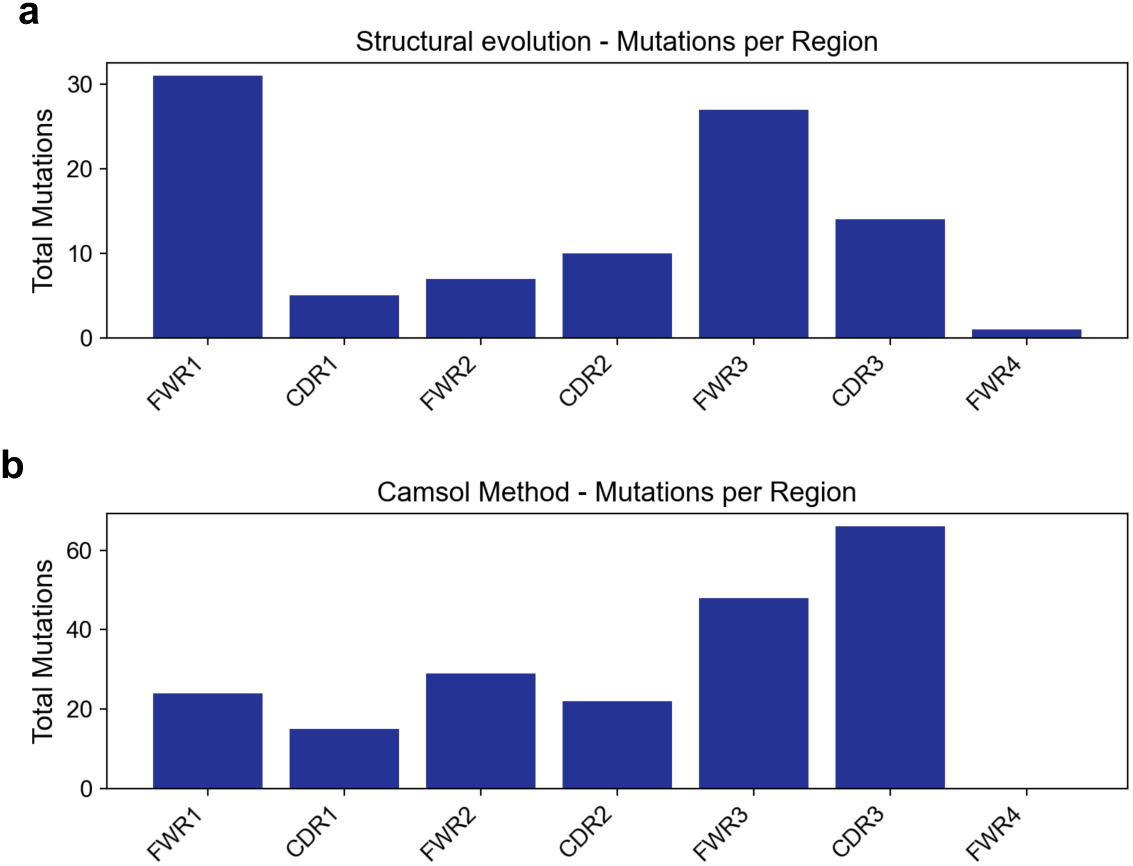
Mutational distribution across V_H_H regions. a) Total number of amino-acid substitutions observed in each V_H_H region (FWR1–4 and CDR1–3) for the structural evolution design set. b) Same analysis for the CamSol design set. Bars report the summed mutation counts per region across all variants, highlighting distinct regional mutational biases between methods (structural evolution concentrates mutations primarily in framework regions, whereas CamSol introduces broader changes with strong enrichment in CDR3 and FWR3.

**Fig. S7.**
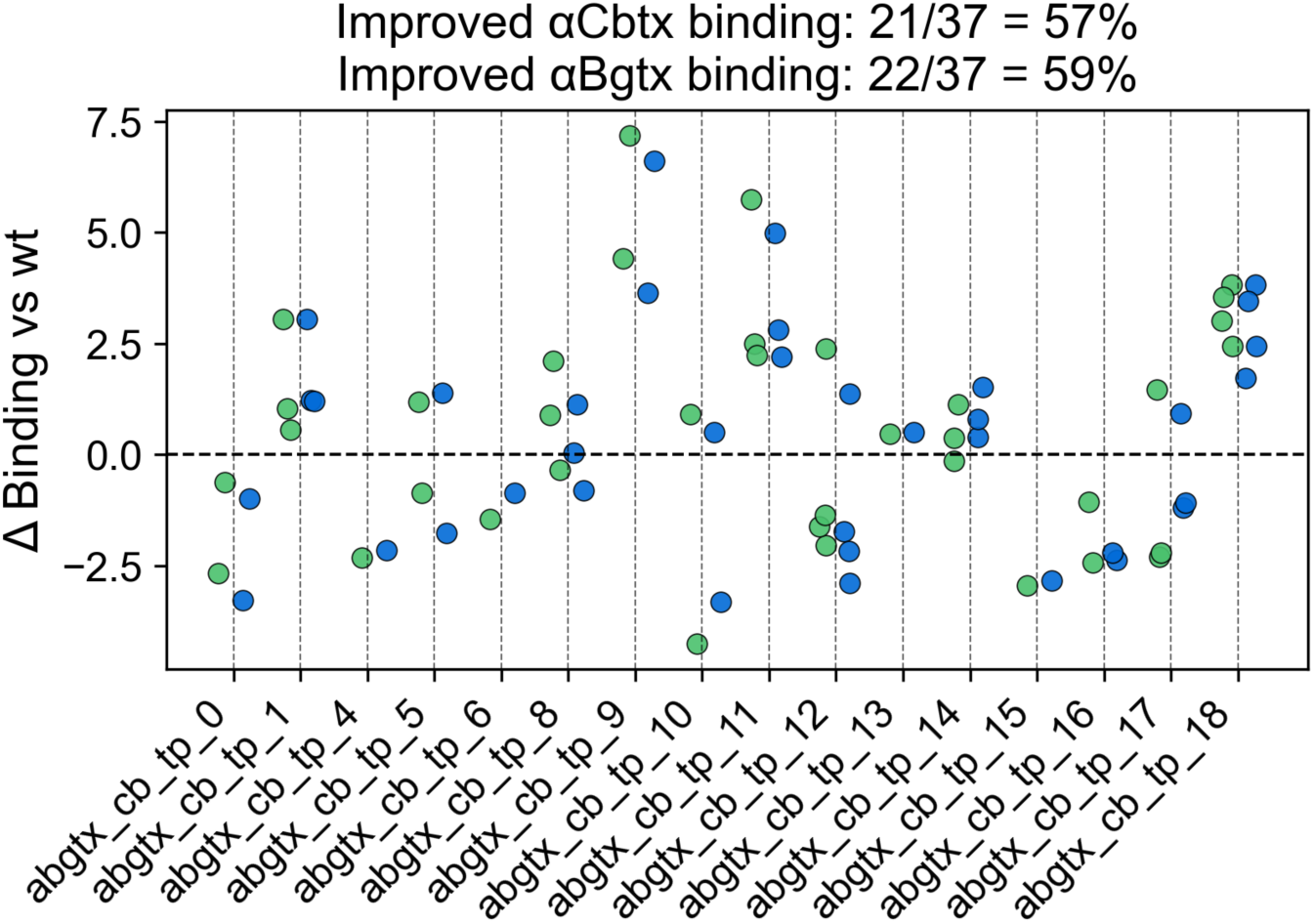
CamSol variants αCbtx and αBgtx binding data. CamSol variants were also tested for binding using the Split-Luciferase binding assay. Shown is the Δ binding of variants relative to the WT. Most variants display comparable or even improved binding signals compared to the WT signal.

**Fig. S8.**
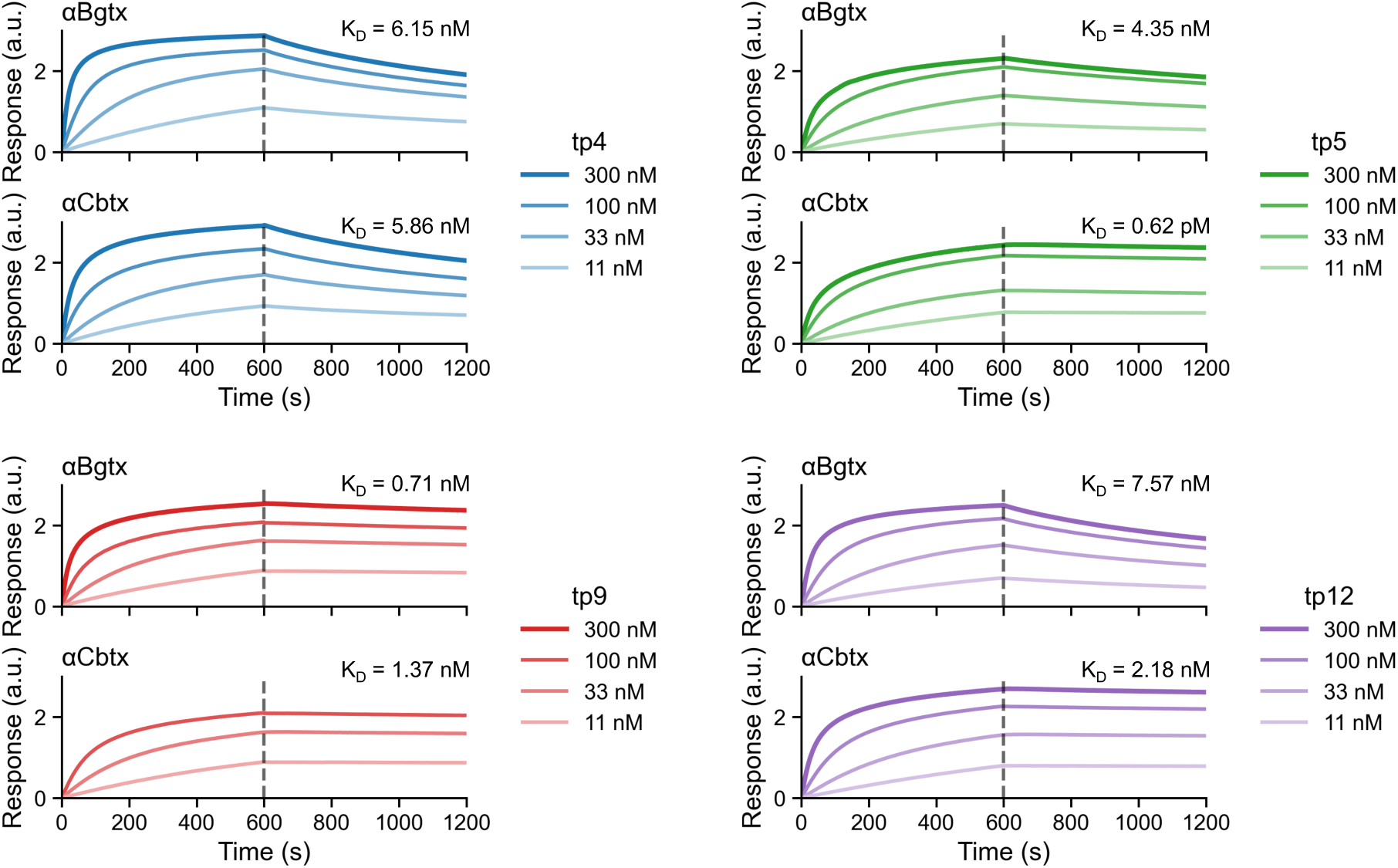
BLI wild-type titrations. Association and dissociation profiles, along with K_d_, are shown for V_H_H tp4, tp5, tp9 and tp12 for αCBTx and αBgtx. All were tested at 4 different concentrations (300 nM, 100 nM, 33 nM and 11 nM).

**Fig. S9.**
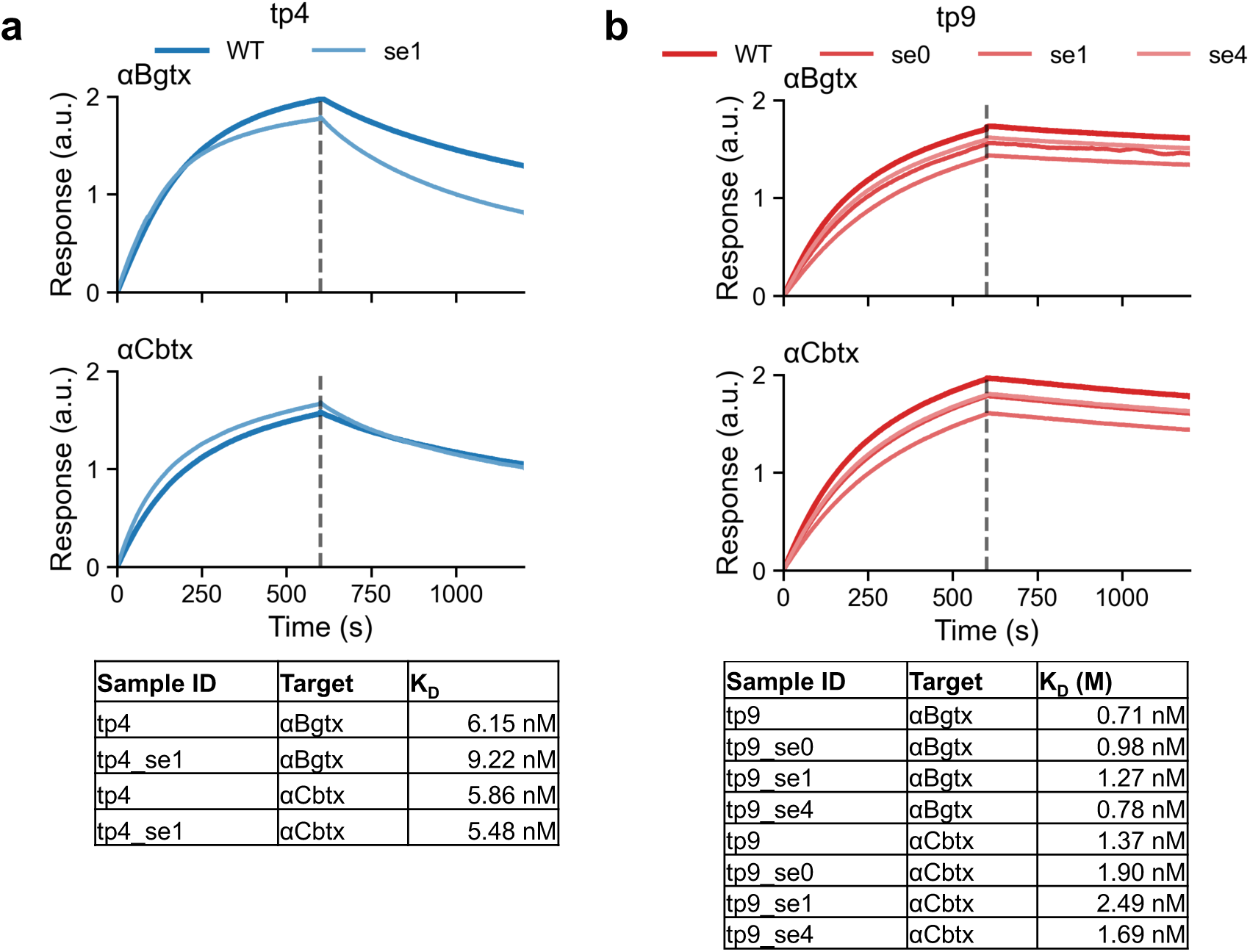
Additional BLI experiments: a) Association and dissociation profiles, along with K_d_, k_on_, and k_dis_, are shown for V_H_H tp4 and one tested variant (tp4_se1). This variant shows comparable affinity for αCbtx, but slightly decreased affinity for αBgtx. b) Affinity data are shown for tp9 and three tested variants. All variants display very similar, though slightly decreased, affinities for both αCbtx and αBgtx.

